# Reduced order modeling and analysis of the human complement system

**DOI:** 10.1101/059386

**Authors:** Adithya Sagar, Wei Dai, Mason Minot, Rachel LeCover, Jeffrey D. Varner

**Affiliations:** Robert Frederick Smith School of Chemical and Biomolecular Engineering, Cornell University, Ithaca, NY, USA

## Abstract

Complement is an important pathway in innate immunity, inflammation, and many disease processes. However, despite its importance, there are few validated mathematical models of complement activation. In this study, we developed an ensemble of experimentally validated reduced order complement models. We combined ordinary differential equations with logical rules to produce a compact yet predictive model of complement activation. The model, which described the lectin and alternative pathways, was an order of magnitude smaller than comparable models in the literature. We estimated an ensemble of model parameters from *in vitro* dynamic measurements of the C3a and C5a complement proteins. Subsequently, we validated the model on unseen C3a and C5a measurements not used for model training. Despite its small size, the model was surprisingly predictive. Global sensitivity and robustness analysis suggested complement was robust to any single therapeutic intervention. Only the simultaneous knockdown of both C3 and C5 consistently reduced C3a and C5a formation from all pathways. Taken together, we developed a validated mathematical model of complement activation that was computationally inexpensive, and could easily be incorporated into pre-existing or new pharmacokinetic models of immune system function. The model described experimental data, and predicted the need for multiple points of therapeutic intervention to fully disrupt complement activation.

## Introduction

Complement is an important pathway in innate immunity. It plays a significant role in inflammation, host defense as well as many disease processes. Complement was discovered in the late 1880s where it was found to ‘complement’ the bactericidal activity of natural antibodies [1]. However, research over the past decade has suggested the importance of complement extends beyond innate immunity. For example, complement contributes to tissue homeostasis [2]. It has also has been linked with several diseases including Alzheimers, Parkinson’s, multiple sclerosis, schizophrenia, rheumatoid arthritis and sepsis [3, 4]. Complement also plays positive and negative roles in cancer; attacking tumor cells with altered surface proteins in some cases, while potentially contributing to tumor growth in others [5, 6]. Lastly, several other important biochemical systems are integrated with complement including the coagulation cascade, the autonomous nervous system and inflammation [6]. Thus, complement is important in a variety of beneficial and potentially harmful functions in the body. Despite its importance, there have been few approved complement specific therapeutics, largely because of safety concerns and challenging pharmacokinetic constraints, however, progress is being made [7].

The complement cascade involves many soluble and cell surface proteins, receptors and regulators [8, 9]. The outputs of complement are the Membrane Attack Complex (MAC), and the inflammatory mediator proteins C3a and C5a. The membrane attack complex, generated during the terminal phase of the response, forms transmembrane channels which disrupt the membrane integrity of targeted cells, leading to cell lysis and death. On the other hand, the C3a and C5a proteins act as a bridge between innate and adaptive immunity, and play an important role in regulating inflammation [5]. Complement activation takes places through three pathways: the classical, the lectin and the alternative pathways. The classical pathway is triggered by antibody recognition of foreign antigens or other pathogens. A multimeric protein complex C1 binds antibody-antigen complexes and undergoes a conformational change, leading to an activated form with proteolytic activity. The activated C1-complex cleaves soluble complement proteins C4 and C2 into C4a, C4b, C2a and C2b, respectively. The C4a and C2b fragments bind to form the C4bC2a protease, also known as the classical pathway C3 convertase (C4bC2a). The lectin pathway is initiated through the binding of L-ficolin or Mannose Binding Lectin (MBL) to carbohydrates on the surfaces of bacterial pathogens. These complexes, in combination with mannose-associated serine proteases 1 and 2 (MASP-1/2), also cleave C4 and C2, leading to additional C4bC2a. Thus, the classical and lectin pathways, initiated by different cues on foreign surfaces, converge at the C4bC2a. On the other hand, the alternative pathway is activated by a ‘tickover’ mechanism in which complement protein C3 is spontaneously hydrolyzed to form an activated intermediate C3w; C3w recruits factor B and factor D, leading to the formation of C3wBb. C3wBb cleaves C3 into C3a and C3b, where the C3b fragment further recruits additional factor B and factor D to form C3bBb, the alternative C3 convertase (AP C3 convertase) [10]. The role of classical and alternative C3 convertases is varied. First, AP C3 convertases mediate signal amplification. AP C3 convertases cleave C3 into C3a and C3b; the C3b fragment is then free to form additional alternative C3 convertases, thereby forming a positive feedback loop. Next, AP/C4bC2as link complement initiation with the terminal phase of the cascade through the formation of C5 convertases. Both classical and alternative C3 convertases can recruit C3b subunits to form the classical pathway C5 convertase (C4bC2aC3b, CP C5 convertase), and the alternative pathway C5 convertase (C3bBbC3b, AP C5 convertase), respectively. Both C5 convertases cleave C5 into the C5a and C5b fragments. The C5b fragment, along with the complement proteins C6, C7, C8 and multiple C9s, form the membrane attack complex. On the other hand, both C3a and C5a are important inflammatory signals involved in several responses [8, 9]. Thus, the complement cascade attacks invading pathogens, while acting as a beacon for adaptive immunity.

The complement cascade is regulated by plasma and host cell surface proteins which balance host safety with effectiveness. The initiation of the classical pathway via complement protein C1 is controlled by the C1 Inhibitor (C1-Inh); C1-Inh irreversibly binds to and deactivates the active subunits of C1, preventing chronic complement activation [11]. Regulation of upstream processes in the lectin and alternative pathways also occurs through the interaction of the C4 binding protein (C4BP) with C4b, and factor H with C3b [12]. Interestingly, both factor H and C4BP are capable of binding their respective targets while in convertase complexes as well. At the host cell surface, membrane cofactor protein (MCP or CD46) can interact with C4b and C3b, which protects the host cell from complement self-activation [13]. Delay accelerating factor (DAF or CD55) also recognizes and dissociates both C3 and C5 convertases on host cell surfaces [14]. More generally the well known inflammation regulator Carboxypeptidase-N has broad activity against the complement proteins C3a, C4a, and C5a, rendering them inactive by cleavage of carboxyl-terminal arginine and lysine residues [15]. Although Carboxypeptidase-N does not directly influence complement activation, it silences the important inflammatory signals produced by complement. Lastly, assembly of the MAC complex itself can be inhibited by vitronectin and clusterin in the plasma, and CD59 at the host surface [16, 17]. Thus, there are many points of control which influence complement across the three activation pathways.

Developing quantitative mathematical models of complement could be crucial to fully understanding its role in the body. Traditionally, complement models have been formulated as systems of linear or non-linear ordinary differential equations (ODEs). For example, Hirayama et al., modeled the classical complement pathway as a system of linear ODEs [18], while Korotaevskiy and co-workers modeled the classical, lectin and alternative pathways as a system of non-linear ODEs [19]. More recently, large mechanistic models of sections of complement have also been proposed. For example, Liu et al., analyzed the formation of the classical and lectin C3 convertases, and the regulatory role of C4BP using a system of 45 non-linear ODEs with 85 parameters [20]. Zewde and co-workers constructed a detailed mechanistic model of the alternative pathway which consisted of 107 ODEs and 74 kinetic parameters and delineated between the fluid, host and pathogen surfaces [17]. However, these previous studies involved large models. The central challenge of complement model identification is the estimation of model parameters from potentially limited experimental measurements. Unlike other important cascades, such as coagulation where there are well developed experimental tools and publicly available data sets, the data for complement is relatively sparse. Data sets with missing or incomplete data, and limited dynamic data also make the identification of large mechanistic complement models difficult. Thus, reduced order approaches which describe the biology of complement using a limited number of species and parameters could be important for pharmacokinetic model development, and for our understanding of the varied role of complement in the body.

## Materials and methods

### Formulation and solution of the complement model equations

We used ordinary differential equations (ODEs) to model the time evolution of complement proteins (*x_i_*) in the reduced order model:

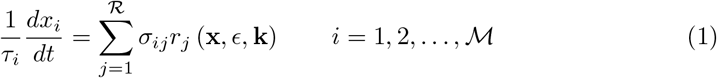

where *R* denotes the number of reactions and *M* denotes the number of proteins in the model. The quantity *τ_i_* denotes a time scale parameter for species *i* which captures unmodeled effects. For the current study, *τ* scaled with the level of initiator (*z*) for C5a and C5b; *τ_i_* = *z/z^∗^* for i = C5a, C5b where *z^∗^* was 1mg/ml, *τ_i_* = 1 for all other species. The quantity *r_j_* (**x**, ∈, **k**) denotes the rate of reaction *j*. Typically, reaction *j* is a non-linear function of biochemical and enzyme species abundance, as well as unknown model parameters **k** (*K ×* 1). The quantity *σ_ij_* denotes the stoichiometric coefficient for species *i* in reaction *j*. If *σ_ij_ >* 0, species *i* is produced by reaction *j*. Conversely, if *σ_ij_ <* 0, species *i* is consumed by reaction *j*, while *σ_ij_* = 0 indicates species *i* is not connected with reaction *j*. Species balances were subject to the initial conditions **x** (*t_o_*) = **x**_*o*_.

Rate processes were written as the product of a kinetic term (*r¯ _j_*) and a control term (*v_j_*) in the complement model. The kinetic term for the formation of C4a, C4b, C2a and C2b, lectin pathway activation, and C3 and C5 convertase activity was given by:

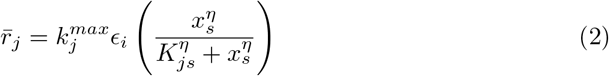

where *k_j_^max^* denotes the maximum rate for reaction *j*, є_*i*_ denotes the abundance of the enzyme catalyzing reaction *j*, *η* denotes a cooperativity parameter, and *K_js_* denotes the saturation constant for species *s* in reaction *j*. We used mass action kinetics to model protein-protein binding interactions within the network:

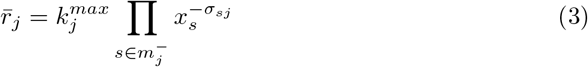

where *k_j_^max^* denotes the maximum rate for reaction *j*, *σ_sj_* denotes the stoichiometric coefficient for species *s* in reaction *j*, and *s* є *m_j_* denotes the set of *reactants* for reaction *j*. We assumed all binding interactions were irreversible.

The control terms 0 ≤ *v_j_* ≤ 1 depended upon the combination of factors which influenced rate process *j*. For each rate, we used a rule-based approach to select from competing control factors. If rate j was influenced by 1,…, *m* factors, we modeled this relationship as *v_j_* = *I_j_* (*f*_1*j*_ (*·*),…, *f_mj_* (*·*)) where 0 *≤ f_ij_* (*·*) *≤* 1 denotes a regulatory transfer function quantifying the influence of factor *i* on rate *j*. The function *I_j_* ( ) is an integration rule which maps the output of regulatory transfer functions into a control variable. Each regulatory transfer function was modeled using a Hill function. In this study, we used *I_j_* є {*min, max*} [21]. If a process has no modifying factors, *v_j_* = 1. The model equations were implemented in Julia and solved using the CVODE routine of the Sundials package [22, 23].

#### Model code repository

The model code and parameter ensemble can be downloaded under an MIT software license from the Varnerlab GitHub repository [24]. The model code, optimization code and analysis routines are maintained in a GitHub repository. The model equations and kinetic rate expressions are encoded in `Balances.jl` which is called by the `SolveBalances.jl` driver function. Note, the user should not directly call `SolveBalances.jl`. Rather, multiple parameter sets can be simulated by calling the driver function from a script. The kinetic and other model parameters are encoded in `DataFile.jl` as a dictionary. The parameters stored in this dictionary can be updated in memory to run different simulations. An example script to simulate the model over the parameter ensemble is encoded in `sample_ensemble.jl`. Plotting routines are encoded in the `PlotLib.jl` library; these routines plot the experimental data contained in the `data` subdirectory. The objective functions, which compute the squared residual between the simulations and experimental measurements, are encoded in the `complement_lib.jl` library. Lastly, we included a set of routines e.g., `Make_Fig_2A.jl` to solve the model equations, and plot the results versus the experimental data for each simulation figure in this study.

### Estimating complement model parameters

A single initial parameter set was estimated using the Dynamic Optimization with Particle Swarms (DOPS) technique [25]. DOPS is a novel hybrid meta-heuristic which combines a multi-swarm particle swarm method with the dynamically dimensioned search approach of Shoemaker and colleagues [26]. DOPS minimized the squared residual between simulated and C3a and C5a measurements with and without zymosan as a single objective. The best fit set estimated by DOPS served as the starting point for multiobjective ensemble generation using Pareto Optimal Ensemble Technique in the Julia programming language (JuPOETs) [27]. JuPOETs is a multiobjective approach which integrates simulated annealing with Pareto optimality to estimate model ensembles on or near the optimal tradeoff surface between competing training objectives. JuPOETs minimized training objectives of the form:

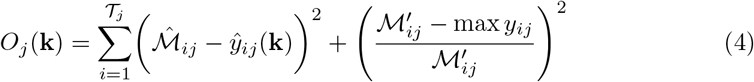

subject to the model equations, initial conditions and parameter bounds *ℒ* ≤ k ≤ *𝒰*. The first term in the objective function measured the shape difference between the simulations and measurements. The symbol *ℳ̂*_*ij*_ denotes a scaled experimental observation (from training set *j*) while the symbol *ŷ*_*ij*_ denotes the scaled simulation output (from training set *j*). The quantity *i* denotes the sampled time-index and 𝒯_*j*_ denotes the number of time points for experiment *j*. The scaled measurement is given by:

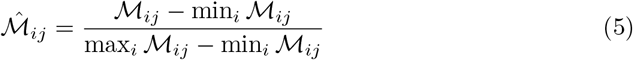

Under this scaling, the lowest measured concentration become zero while the highest equaled one, where a similar scaling was defined for the simulation output. The second-term in the objective function quantified the absolute error in the estimated concentration scale, where the absolute measured concentration (denoted by *ℳ*ʹ_*ij*_) was compared with the largest simulated value. In this study, we minimized two training objectives, the total C3a and C5a residual w/o zymosan (O_1_) and the total C3a and C5a residual for 1 mg/ml zymosan (O_2_). JuPOETs identified an ensemble of N = 2100 parameter sets which were used for model simulations and uncertainty quantification subsequently. JuPOETs is open source, available under an MIT software license. The JuPOETs source code is freely available from the JuPOETs GitHub repository [28]. The objective functions (and experimental data) used in this study are available from the Varnerlab GitHub repository [24].

The simulation and prediction performance of the complement model was measured using the Akaike information criterion (AIC) [29]. In this study, we implemented the AIC as:

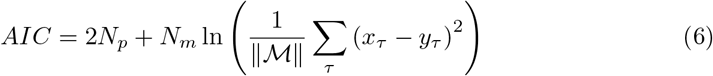

where *N_p_, N_m_* denotes the number of parameters, and the number of experimental measurements, respectively. The summation term in Eq. (6) denotes the residual between the model simulation (*x*) and experimental measurements (*y*), where the residual is normalized by the scale of the experimental data (∥ℳ∥). We compared the AIC for the model parameters estimated in this study, with a random parameter control generated to have a similar order of magnitude. The mean and standard deviation of the AIC was calculated over the parameter ensemble and the random parameter control were reported in this study.

#### Global sensitivity analysis

We conducted global sensitivity analysis to estimate which parameters and species controlled the performance of the reduced order model. We computed the total variance-based sensitivity index of each parameter relative to the training residual for the C3a/C5a alternative and C3a/C5a lectin objectives using the Sobol method [30]. The sampling bounds for each parameter were established from the minimum and maximum value for that parameter in the parameter ensemble. We used the sampling method of Saltelli *et al*. to compute a family of *N* (2*d* + 2) parameter sets which obeyed our parameter ranges, where *N* was the number of trials per parameters, and *d* was the number of parameters in the model [31]. In our case, *N* = 400 and *d* = 28, so the total sensitivity indices were computed using 23,200 model evaluations. The variance-based sensitivity analysis was conducted using the SALib module encoded in the Python programming language [32].

#### Pairwise sensitivity analysis and clustering

We perturbed each pair of model parameters by 10% of their nominal value, and then calculated the euclidian distance between the perturbed and nominal system states for physiological conditions. We repeated this calculation for each member of the parameter ensemble, and calculated the mean differences between the perturbed and nominal states. We then clustered the resulting `log10` transformed mean distances using the `Clustergram` routine in MATLAB (The Mathworks, Natick MA). We considered three clusters, high, medium and low displacement.

#### Robustness analysis

Robustness coefficients quantify the response of a marker to a structural or operational perturbation to the network architecture. Robustness coefficients were calculated as shown previously [33]. Log-transformed robustness coefficients denoted by α̂ (*i, j, t_o_, t_f_*) were defined as:

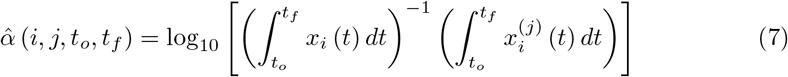

Here, *t_o_* and *t_f_* denote the initial and final simulation time, while *i* and *j* denote the indices for the marker and the perturbation, respectively. A value of *α*ˆ (*i, j, t_o_, t_f_*) *>* 0, indicates increased marker abundance, while *α*ˆ (*i, j, t_o_, t_f_*) *<* 0 indicates decreased marker abundance following perturbation *j*. If *α*ˆ (*i, j, t_o_, t_f_*) ~ 0, perturbation *j* did not influence the abundance of marker *i*. In this study, we perturbed the initial condition of C3 or C5 or a combination of C3 and C5 by 50%, 90% and 99% and measured the area under the curve (AUC) of C3a or C5a with and without lectin initiator. We computed the robustness coefficients for a subset of the parameter ensemble (N = 65) and reported the mean robustness value.

## Results

In this study, we estimated an ensemble of experimentally validated reduced order complement models using multiobjective optimization. The modeling approach combined ordinary differential equations with logical rules to produce a complement model with a limited number of equations and parameters. The reduced order model, which described the lectin and alternative pathways, consisted of 18 differential equations with 28 parameters. Thus, the model was an order of magnitude smaller than comparable models in the literature. We estimated an ensemble of model parameters from *in vitro* time series measurements of the C3a and C5a complement proteins. Subsequently, we validated the model on unseen C3a and C5a measurements not used for model training. Despite its size, the model was surprisingly predictive. After validation, we performed global sensitivity and robustness analysis to estimate which parameters and species controlled model performance. Sensitivity analysis suggested CP C3 and C5 convertase parameters were critical, while robustness analyses suggested complement was robust to any single therapeutic intervention; only the knockdown of both C3 and C5 consistently reduced C3a and C5a formation for all cases. Taken together, we developed a reduced order complement model that was computationally inexpensive, and could easily be incorporated into pre-existing or new pharmacokinetic models of immune system function. The model described experimental data, and predicted the need for multiple points of intervention to disrupt complement activation.

### Reduced order complement network

The complement model described the initiation of the alternative and lectin pathways, and the downstream integration of the lectin and classical pathways (Fig. 1). A trigger event initiated the lectin pathway (encoded as a logical rule), which activated the cleavage of C2 and C4 into C2a, C2b, C4a and C4b, respectively. In this study, we did not explicitly model classical pathway activation mediated by the C1 protein. Instead we described the integration of the lectin and classical pathways at the level of C2 and C4 activation and the formation of the Classical Pathway (CP) C3 convertase (C4aC2b). However, in future studies classical pathway activation could be easily described by simply adding additional terms to the C2/C4 and C2a, C2b, C4a and C4b balances, leaving all other components largely unchanged. Classical Pathway (CP) C3 convertase (C4aC2b) then catalyzed the cleavage of C3 into C3a and C3b. The alternative pathway was initiated through the spontaneous hydrolysis of C3 into C3a and C3b. The C3b fragments generated by hydrolysis (or by C4bC2a) could then form the alternative pathway (AP) C3 convertase (C3bBb). However, in this study we considered lumped alternative pathway activation; we did not consider C3w, nor the formation of the initial alternative C3 convertase (C3wBb). Rather, we assumed C3w was equivalent to C3b and only modeled the formation of the main AP C3 convertase. Both the CP and AP C3 convertases catalyzed the cleavage of C3 into C3a and C3b. A second C3b fragment could then bind with either the CP or AP C3 convertase to form the CP or AP C5 convertase (C4bC2aC3b or C3bBbC3b). Both C5 convertases catalyzed the cleavage of C5 into the C5a and C5b fragments. In this study, we simplified the model by assuming both factor B and factor D were in excess. However, we did explicitly account for the action of two other control proteins, factor H and C4BP. Lastly, we did not consider MAC formation, instead we stopped at C5a and C5b. Lectin pathway activation, and C3/C5 convertase activity were modeled using a combination of saturation kinetics and non-linear transfer functions, which resulted in a significant size reduction of the model, while maintaining performance. Binding interactions were modeled using mass-action kinetics, where we assumed all binding was irreversible. Thus, while the reduced order complement model encoded significant biology, it was highly compact consisting of only 18 differential equations and 28 model parameters. Next, we estimated an ensemble of model parameters from time series measurements of the C3a and C5a complement proteins.

**Figure 1.**
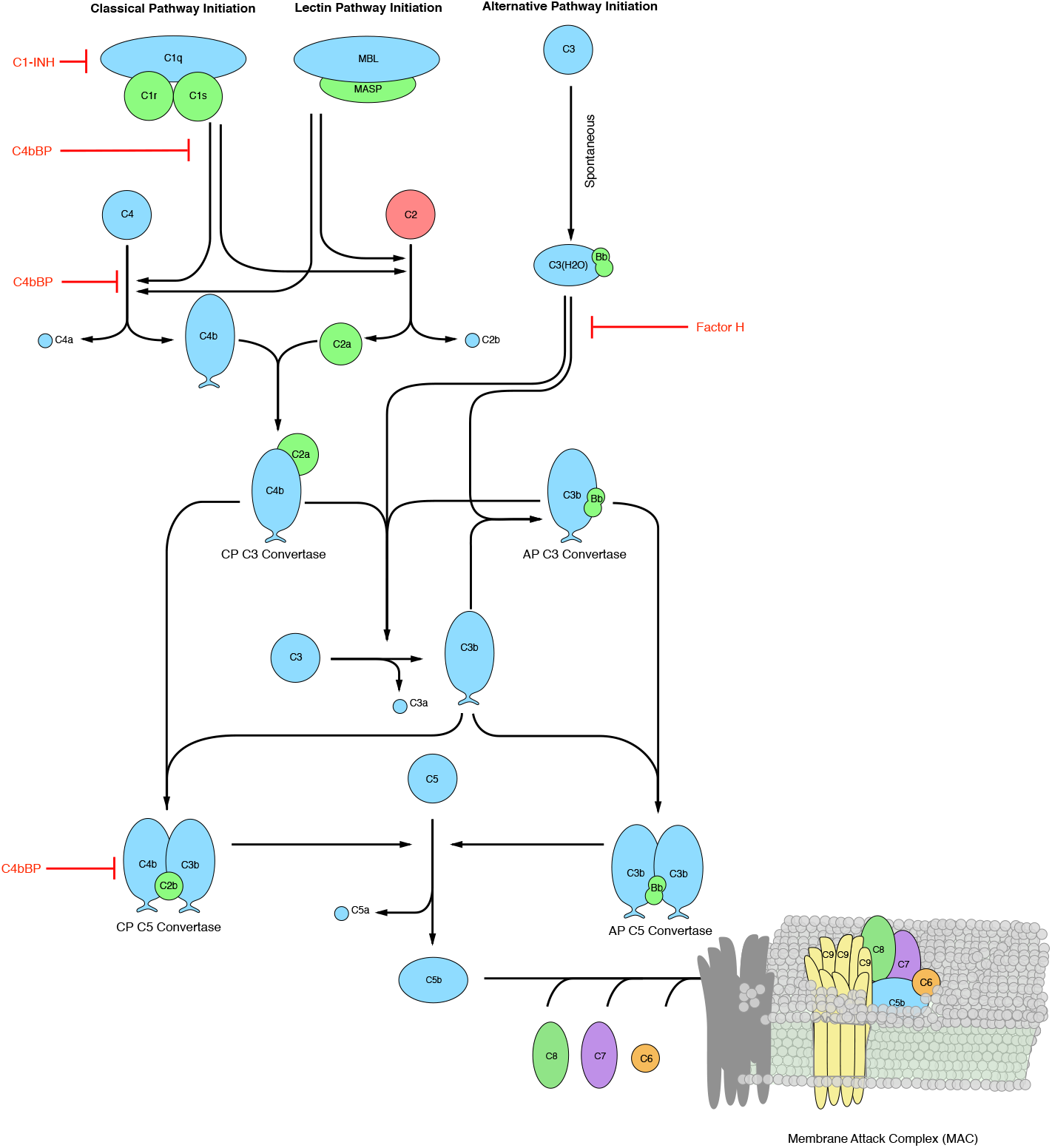
Simplified schematic of the human complement system. The complement cascade is activated through three pathways: the classical, the lectin, and the alternative pathways. Complement initiation results in the formation of classical or alternative C3 convertases, which amplify the initial complement response and signal to the adaptive immune system by cleaving C3 into C3a and C3b. C3 convertases further react to form C5 convertases which catalyze the cleavage of the C5 complement protein to C5a and C5b. C5b is critical to the formation of the membrane attack complex (MAC), while C5a recruits an adaptive immune response.

### Estimating an ensemble of reduced order complement models

A critical challenge for the development of any dynamic model is the estimation of model parameters. We estimated an ensemble of complement model parameters using *in vitro* time-series data sets generated with and without zymosan A, a complement pathway activator. The data used for model training was taken from the study of Morad et al. [34] and is given in Table T2. In this study, we assumed zymosan A differentially activated the lectin pathway, consistent with the study of Morad et al [34] and several previous studies [35–37]. However, the role of zymosan may be more complex, as it may activate all three complement pathways under certain conditions. The residual between model simulations and experimental measurements was minimized using the Pareto Optimal Ensemble Technique (JuPOETs) [38]. starting from a initial guess generated by the dynamic optimization with particle swarms (DOPS) routine. Unless otherwise specified, all initial conditions were assumed to be at their mean physiological values. While we had significant training data, the parameter estimation problem was underdetermined (we were not able to uniquely determine model parameters). Thus, instead of using the best-fit yet uncertain parameter set, we estimated an ensemble of probable parameter sets to quantify model uncertainty (N = 2100, see materials and methods). The complement model ensemble captured the behavior of both the alternative and lectin pathways (Fig. 2). To estimate alternative pathway model parameters, we used C3a and C5a measurements in the absence of zymosan (Fig. 2A and B). On the other hand, lectin pathway parameters were estimated from C3a and C5a measurements in the presence of 1mg/ml zymosan (Fig. 2C and D). The reduced order model reproduced a panel of alternative and lectin pathway data sets in the neighborhood of physiological factor and inhibitor concentrations. The model fit for parameter sets estimated by JuPOETs, quantified by the Akaike information criterion (AIC), was statistically significantly different than a random parameter control for each case at a 95% confidence level. However, it was unclear whether the reduced order model could predict new data, without updating the model parameters. To address this question, we fixed the model parameters and simulated data sets not used for model training.

**Figure 2.**
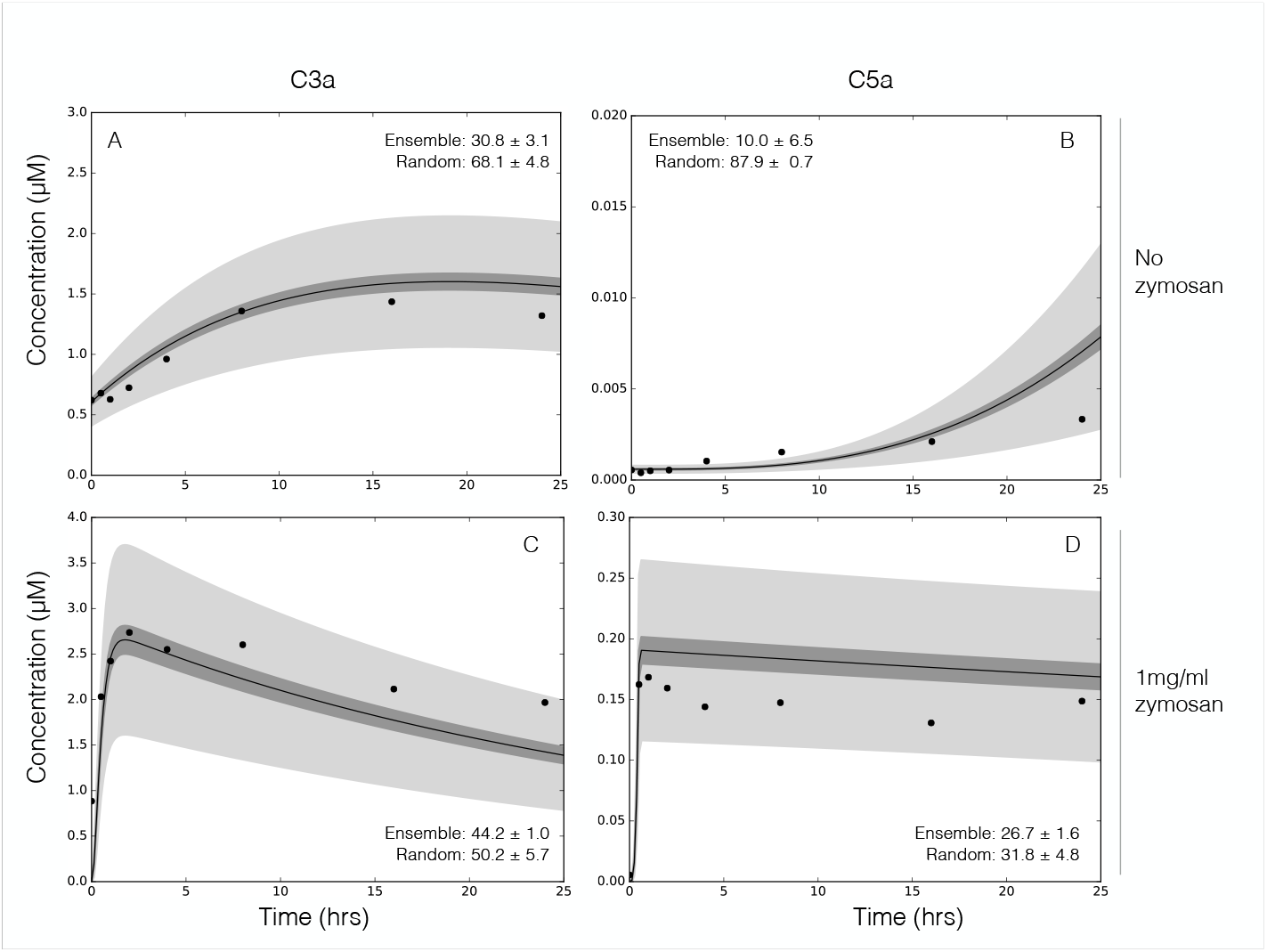
Reduced order complement model training. An ensemble of model parameters was estimated using multiobjective optimization from dynamic C3a and C5a measurements with and without zymosan. The model was trained using C3a and C5a measurements in the absence of zymosan (**A**–**B**) or in the presence of 1 mg/ml zymosan (**C**–**D**). The solid black lines show the simulated mean value of C3a or C5a for the ensemble, while the dark shaded region denotes the 99% confidence interval of mean. The light shaded region denotes the 99% confidence interval of the simulated C3a and C5a concentration. The experimental training data (points) was taken from Morad et al [34]. All initial conditions not specified by the experimental condition were assumed to be at zero or their physiological serum levels unless otherwise noted (Table T1).

We tested the predictive power of the reduced order complement model with data not used during model training (Fig. 3). The data used for model validation was taken from the study of Morad et al. [34] and is given in Table T2. Six validation cases were considered, three for C3a and C5a each, respectively. Similar to model training, we compared the AIC for each prediction case to a randomized parameter family. All model parameters and initial conditions were fixed for the validation simulations (with the exception of zymosan, and other experimentally mandated changes). The ensemble of reduced order models predicted the qualitative dynamics of C3a formation (Fig. 3, top), and C5a formation (Fig. 3, bottom) at three inducer concentrations. For each training case, the AIC was statistically significantly different than the random parameter control for a 95% confidence level. The rate of C3a formation and C3a peak time were directly proportional to initiator dose. Similarly, the C5a plateau and rate of formation were also directly proportional to initiator dose, with the lag time being indirectly proportional to initiator exposure for both C3a and C5a. However, there were shortcomings with model performance. First, while the overall C3a trend was captured (within the 99% confidence interval), the C3a dynamics were too fast with the exception of the low dose case. We believe the C3a time scale was related to our choice of training data, how we modeled the tickover mechanism, and factor B and D limitation. We trained the model using either no or 1 mg/ml zymosan, but predicted cases in a different initiator range; comparing training to prediction, the model performance e.g., the shape of the C3a trajectory was biased towards either high or very low initiator doses. Next, tickover was modeled as a first-order generation processes where C3wBb formation and activity was lumped into the AP C3 convertase. Thus, we skipped an important upstream step which could influence AP C3 convertase formation by attenuating the rate C3 cleavage into C3a and C3b. We also assumed both factor B and factor D were not limiting, thereby artificially accelerating the rate of AP C3 convertase formation. The C5a predictions followed a similar trend as C3a; we captured the long-time C5a behavior but over predicted the time scale of C5 cleavage. However, because the C5a time scale depends strongly upon C3 convertase formation, we can likely correct the C5 issues by fixing the rate of C3 cleavage. Despite these shortcomings, we qualitatively predicted experimental measurements not used for model training typically within the 99% confidence of the ensemble, for three inducer levels. Next, we used global sensitivity and robustness analysis to determine which parameters and species controlled the performance of the complement model.

**Figure 3.**
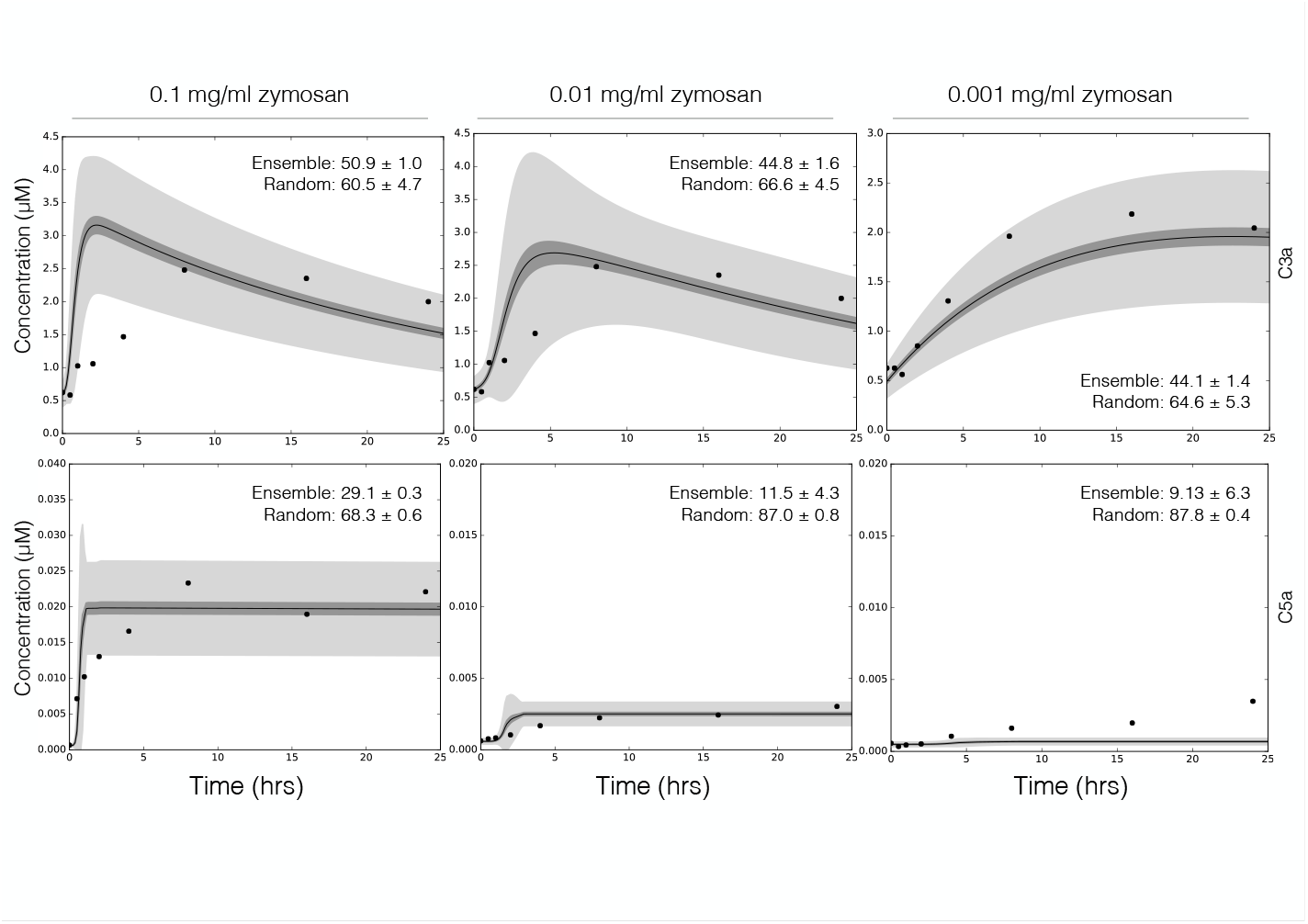
Reduced order complement model predictions. Simulations of C3a and C5a generated using 0.1 mg/ml, 0.01 mg/ml, and 0.001 mg/ml zymosan were compared with the corresponding experimental measurements. The solid black lines show the simulated mean value of C3a or C5a for the ensemble, while the dark shaded region denotes the 99% confidence interval of mean. The light shaded region denotes the 99% confidence interval of the simulated C3a and C5a concentration. The experimental validation data (points) was taken from Morad et al [34]. All initial conditions not specified by the experimental condition were assumed to be at zero or their physiological serum levels unless otherwise noted (Table T1).

### Global analysis of the reduced order complement model

We conducted sensitivity analysis to estimate which parameters controlled the performance of the reduced order complement model. We calculated the total sensitivity of the C3a and C5a residual to changes in model parameters with and without zymosan (Fig. 4). In the absence of zymosan (where only the alternative pathway is active), the most sensitive parameter was the rate constant governing the assembly of the AP C3 convertase, as well as the rate constant controlling basal C3b formation via the tickover mechanism. The C5a trajectory was sensitive to the AP C5 convertase kinetic parameters (Fig. 4A). Interestingly, neither the rate nor the saturation constant governing AP C3 convertase activity were sensitive in the absence of zymosan. Thus, C3a formation in the alternative pathway was more heavily influenced by the spontaneous hydrolysis of C3, rather than AP C3 convertase activity, in the absence of zymosan. In the presence of zymosan, the C3a residual was controlled by the formation and activity of the C4bC2a, as well as tickover and degradation parameters. On the other hand, the C5a residual was controlled by the formation and activity of CP C5 convertase, and tickover C3b formation in the presence of zymosan (Fig. 4B). The lectin initiation parameters were sensitive, but to a lesser extent than CP convertase kinetic parameters and tickover C3b formation. Thus, sensitivity analysis suggested that CP C3/C5 convertase formation and activity dominated in the presence of zymosan, but tickover parameters and AP C5 convertase were more important without initiator. AP C3 convertase assembly was important, but its activity was not. Next, we compared the sensitivity results to current therapeutic approaches; pathways involving sensitive parameters have been targeted for clinical intervention (Fig. 4C). In particular, the sensitivity analysis suggested AP/CP C5 convertase inhibitors, or interventions aimed at attenuating C3 or C5 would most strongly influence complement performance. Thus, there was at least a qualitative overlap between sensitivity and the potential of biochemical efficacy. However, total sensitivity coefficients quantify how simultaneous changes in many parameters e.g., rate or saturation constants affect model performance (in this case model fit). To better understand the role of each parameter, and parameter combination, we explored how finite changes in parameter combinations influenced model performance.

**Figure 4.**
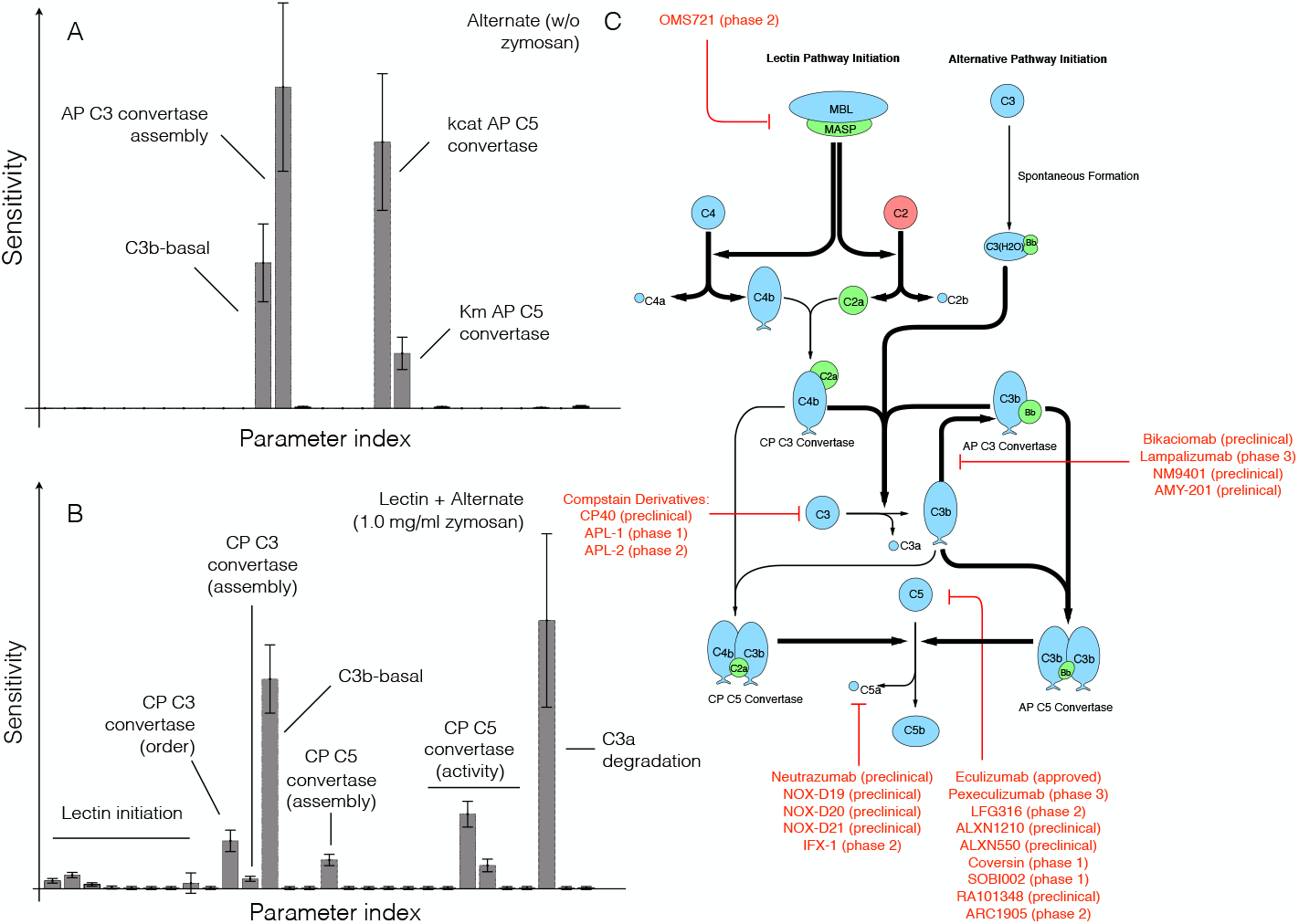
Global sensitivity analysis of the reduced order complement model. Sensitivity analysis was conducted on the two objectives used for model training. **A**: Sensitivity of the C3a and C5a residual w/o zymosan. **B**: Sensitivity of the C3a and C5a residual with 1 mg/ml zymosan. The bars denote the mean total sensitivity index for each parameter, while the error bars denote the 95% confidence interval. **C**: Pathways controlled by the sensitivity parameters. Bold black lines indicate the pathway involves one or more sensitive parameters, while the red lines show current therapeutics targets. Current complement therapeutics were taken from the review of Morgan and Harris [39].

Pairwise parameter perturbations identified crosstalk within the complement model (Fig. 5). We perturbed each pairwise parameter combination by 10%, and calculated the distance between the perturbed and nominal state for each parameter set in the ensemble. We then clustered the mean response of each parameter combination based upon the euclidian distance between the perturbed and nominal states into low (green), medium (red) and high (blue) response clusters. A low response (white) meant the parameter perturbations did not significantly change the system state compared with the nominal case. Four of the 28 parameters (or approximately 14% of the overall model parameters) were in the high response cluster (Fig. 5, blue cluster). These parameters included the rate constant controlling the basal formation of C3b (#12), C3a degradation (#26) as well as the catalytic rate constant governing C4bC2a activity (#22). The only C5 related parameter in the high response group was the rate constant controlling the formation of CP C5 convertase (#15). Approximately, 36%, or 10 of the 28 model parameters, were clustered in the medium impact cluster (Fig. 5, red cluster). Three parameters (#10, #1, #27) were especially important in this cluster; The reaction order governing C4bC2a activity was important (#10), along with the rate constant controlling C4a and C4b formation from C4 in the lectin initiation pathway (#1), and the constant controlling the inhibitory action of C4BP (#27). Lastly, 50% of the model parameters were clustered in the low response cluster (Fig. 5, green cluster). Many of these parameters influenced complement activation; for example, parameter #23 (the C4bC2a saturation constant) was important, just not to the extent of other model parameters. Pairwise synergistic interactions between parameters were also identified. For example, in the high impact cluster, three synergistic relationships were identified, a single positive and two negative cases. Parameters #12 (rate constant governing basal C3b formation) and #15 (formation of CP C5 convertase) acted synergistically to increase the system response. On the other hand, simultaneously changing parameters #12 and #22 or #15 and #26 decreased the system response relative to a single perturbation. However, the most striking examples of synergy occurred in the medium impact cluster; for example, simultaneously increasing parameters #13 (rate constant governing AP C3 convertase formation) and #19 (saturation constant governing AP C5 convertase activity) significantly changed the model state. Changes in parameter #3 (rate constant governing C2a and C2b formation from C2) showed both positive and negative synergistic effects depending upon the other parameter that was perturbed. Taken together, sensitivity coefficients quantified how changes in parameters or parameter combinations affected model performance. However, individual parameters e.g., rate or saturation constants are not easily druggable. To more closely simulate a clinical intervention e.g., administration of anti-complement inhibitors, we performed knock-down analysis on the initial values of C3 and C5 in the absence and presence of flow.

**Figure 5.**
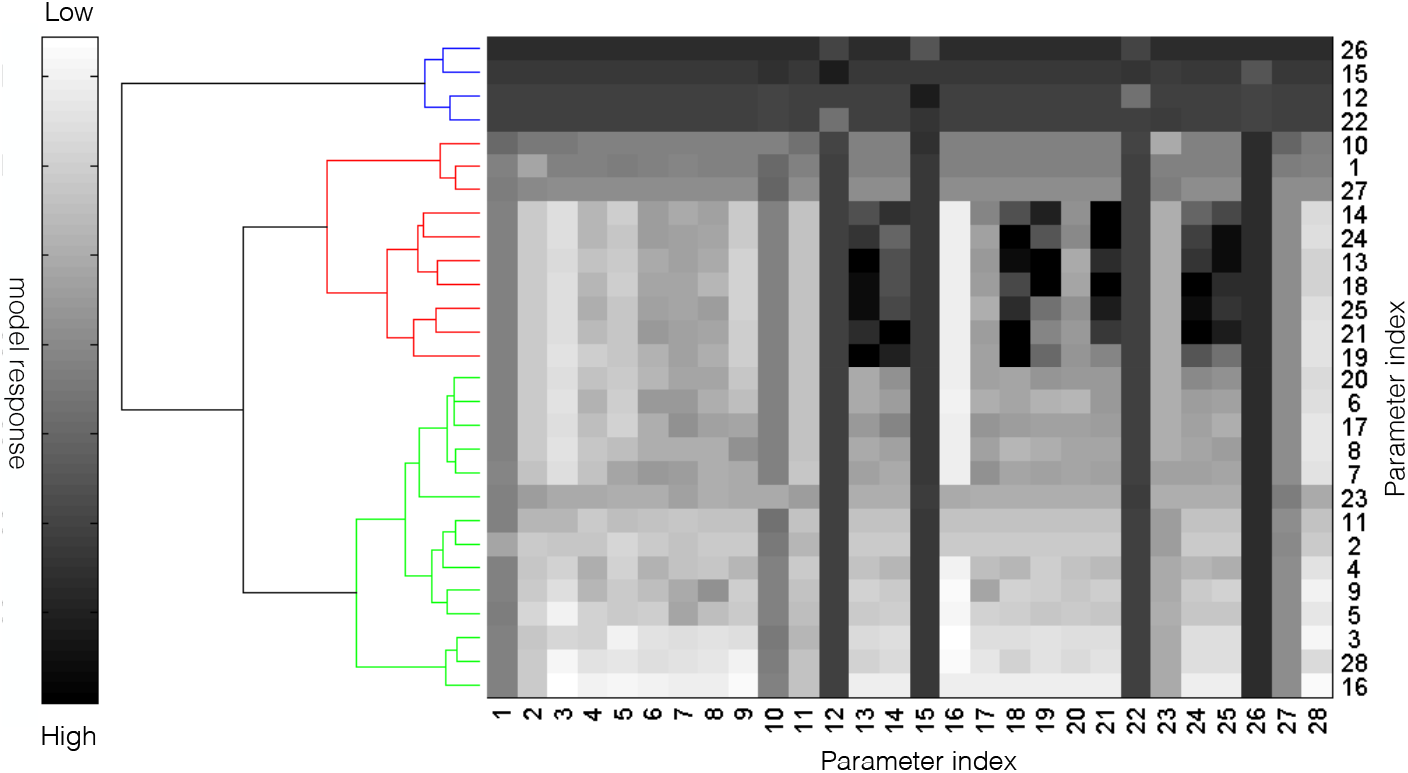
Pairwise sensitivity and clustering of complement model parameters in the presence of 1 mg/ml zymosan. The response of the complement model was calculated for each parameter combination following a 10% increase in parameter combinations in the presence of 1 mg/ml zymosan. The model parameters were clustered into high (blue), medium (red) and low (green) response clusters based upon the euclidian distance between the perturbed and nominal system state (no perturbation).

Knock-down analysis in the absence of flow suggested there was no single intervention that inhibited complement activation in the presence of both initiation pathways (Fig. 6). Robustness coefficients quantify the response of a protein to a macroscopic structural or operational perturbation to a biochemical network. Here, we computed how the C3a and C5a trajectories responded to a decrease in the initial abundance of C3 and/or C5 with and without lectin initiator. We simulated the addition of different doses of anti-complement inhibitor cocktails by decreasing the initial concentration of C3, C5 or the combination of C3 and C5 by 50%, 90% and 99%. This would be conceptually analogous to the administration of a C3 inhibitor e.g., Compstatin alone or combination with Eculizumab (Fig. 4C). The response of the complement model to different knock-down magnitudes was non-linear; a 90% knock-down had an order of magnitude more impact than a 50% knock-down. As expected, a C5 knockdown had no effect on C3a formation for either the alternative (Fig. 6A) or lectin pathways (Fig. 6B). However, C3a and to a greater extent C5a abundance decreased with decreasing C3 concentration in the alternative pathway (Fig. 6A). This agreed with the sensitivity results; changes in AP C3-convertase formation affected the downstream dynamics of C5a formation. Thus, if we only considered the alternative pathway, C3 alone could be a reasonable target, especially given that C5a formation was surprisingly robust to C5 levels in the alternative pathway. Yet, when both pathways were activated, C5a levels were robust to the initial C3 concentration (Fig. 6B); even 1% of the nominal C3 was able to generate enough AP/CP C5 convertase to maintain C5a formation. Thus, the only reliable intervention that consistently reduced both C3a and C5a formation for all cases was a knockdown of both C3 and C5. For example, a 90% decrease of both C3 and C5 reduced the formation of C5a by an order of magnitude, while C3a was reduced to a lesser extent (Fig. 6B). Taken together, these results suggested that both C3 and C5 inhibitors must be administered in the presence of both initiation pathways.

**Figure 6.**
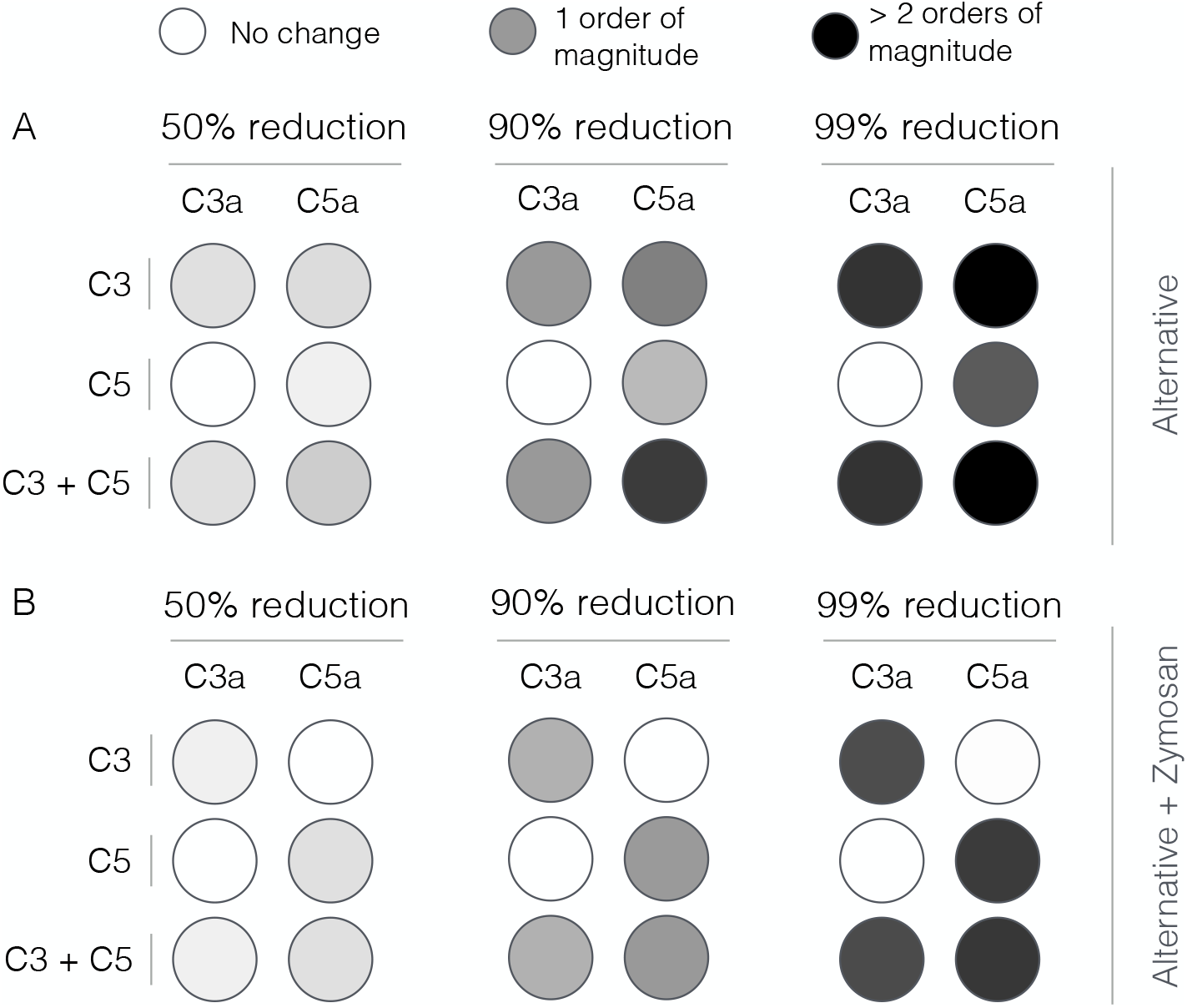
Robustness analysis of the complement model. Robustness coefficients were calculated for a 50%, 90% and 99% reduction in C3, C5, or C3 and C5 initial conditions. **A**: Mean robustness index for C3a and C5a generated in the absence of zymosan. **B**: Mean robustness index for C3a and C5a generated in the presence of 1 mg/ml zymosan. The color describes the degree of reduction of C3a or C5a following the network perturbation. Robustness coefficients were calculated using all parameter sets with Pareto rank less than five (N = 65). Mean robustness values were reported.

## Discussion

In this study, we estimated an ensemble of experimentally validated reduced order complement models using multiobjective optimization. The modeling approach combined ordinary differential equations with logical rules to produce a complement model with a limited number of equations and parameters. The reduced order model, which described the lectin and alternative pathways, consisted of 18 differential equations with 28 parameters. Thus, the model was an order of magnitude smaller than comparable mathematical models in the literature. We estimated an ensemble of model parameters from *in vitro* time series measurements of the C3a and C5a complement proteins. Subsequently, we validated the model on unseen C3a and C5a measurements that were not used for model training. Despite its small size, the model was surprisingly predictive. After validation, we performed global sensitivity and robustness analysis to estimate which parameters and species controlled model performance. These analyses suggested complement was robust to any single therapeutic intervention. The only intervention that consistently reduced C3a and C5a formation for all cases was a knockdown of both C3 and C5. Taken together, we developed a reduced order complement model that was computationally inexpensive, and could easily be incorporated into pre-existing or new pharmacokinetic models of immune system function. The model described experimental data, and predicted the need for multiple points of intervention to disrupt complement activation.

There has been a paucity of validated mathematical models of complement pathway activation. To our knowledge, this study is one of the first complement models that combined multiple initiation pathways with experimental validation of important complement products like C5a. However, there have been several theoretical models of components of the cascade in the literature. Liu and co-workers modeled the formation of C3a through the classical pathway using 45 non-linear ODEs [20]. In contrast, in this study we modeled lectin mediated C3a formation using only five ODEs. Though we did not model all the initiation interactions in detail, especially the cross-talk between the lectin and classical pathways, we successfully captured C3a dynamics with respect to different concentrations of lectin initiators. The model also captured the dynamics of C3a and C5a formed from the alternative pathway using only seven ODEs. The reduced order model predictions of C5a were qualitatively similar to the theoretical complement model of Zewde et al., which involved over 100 ODEs [17]. However, we found that the C3a produced in the alternative pathway was nearly three orders of magnitude greater than the C5a generated. While this was in agreement with the experimental data [34], it differed from the theoretical predictions made by Zewde et al., who showed C3a was eight orders of magnitude greater than the C5a concentration [17]. In our model, the time profile of both C3a and C5a generated changed with respect to the quantity of zymosan (the lectin pathway initiator). In particular, the C3a peak time was directly proportional to initiator, while the lag phase for generation was inversely proportional to the initiator concentration. Korotaevskiy et al. showed a similar trend using a theoretical model of complement, albeit for much shorter time scales [19]. Thus, the reduced order complement model performed at least as well as existing larger mechanistic models, despite being significantly smaller.

Global analysis of the complement model suggested potentially important therapeutic targets. Complement malfunctions are implicated in a spectrum of diseases, however the development of complement specific therapeutics has been challenging [3, 39]. Previously, we have shown that mathematical modeling and analysis can be useful tools to estimate therapeutically important mechanisms [40–43]. In this study, we analyzed a validated ensemble of reduced order complement models to better understand the strengths and weaknesses of the cascade. In the presence of an initiator, C3a and C5a formation was sensitive to CP C3/C5 convertase assembly and activity, and to a lesser extent lectin initiation parameters. Formation of the CP convertases can be inhibited by targeting upstream protease complexes like MASP-1,2 from the lectin pathway (or C1r, C1s from classical pathway). For example, Omeros, a protease inhibitor that targets the MASP-2 complex, has been shown to inhibit the formation of downstream convertases [44]. Lampalizumab and Bikaciomab, which target factor B and factor D respectively, or naturally occurring proteins such as Cobra Venom Factor (CVF), an analogue of C3b, could also attenuate AP convertase formation [45–47]. Removing supporting molecules could also destabilize the convertases. For example, Novelmed Therapeutics developed the antibody, NM9401 against propedin, a small protein that stabilizes alternative C3 convertase [48]. Lastly, convertase catalytic activity could be attenuated using small molecule protease inhibitors. All of these approaches are consistent with the results of the sensitivity analysis. On the other hand, robustness analysis suggested C3a and C5a generation could only be significantly attenuated by modulating the free levels of C3 and C5. The most commonly used anti-complement drug Eculizumab, targets the C5 protein [39]. Several other antibodies targeting C5 are also being developed; for example, LFG316 targets C5 in Age-Related Macular Degeneration [49], while Mubodina is used to treat Atypical Hemolytic-Uremic Syndrome (aHUS) [50]. Other agents such as Coversin [51] or the aptamer Zimura [52] could also be used to knockdown C5. The peptide inhibitor Compstatin and its derivatives are promising approaches for the inhibition of C3 [53]. However, while the knockdown of C3 and C5 affect C3a and C5a levels downstream, the abundance, turnover rate and population variation of these proteins make them difficult targets [54, 55]. For example, the eculizumab dosage must be significantly adjusted during the course of treatment for aHUS [56]. This suggests that therapeutic targets estimated using whole-body models which incorporate pharmacokinetic factors in combination with biochemistry may give higher fidelity predictions of patient response to therapeutic intervention than static biochemical network models.

Despite its importance, there have been few approved complement specific therapeutics because of safety and pharmacokinetic constraints [7]. A validated complement model, in combination with personalized pharmacokinetic models of immune system function, could be an important development for the field. The integration of effective models of complement and other important cascades in the blood, for example the coagulation cascade, with physiologically based pharmacokinetic models is an exciting opportunity. Physiologically based pharmacokinetic models, composed of a collection of organ models (of variable complexity) interconnected by a circulatory system, describe the physical disposition of blood constituents within the body [57]. PBPK models, originally developed to predict the tissue distribution of therapeutic agents or toxins [58], have become an accepted pharmacokinetic tool [59, 60]. PBPK models readily integrate potentially important clinical information such as the demographic and physical characteristics of patients, dissolved oxygen levels, blood flow rates to well- and poorly-perfused organs, pulmonary function and pharmacokinetic factors etc, with blood biochemistry. Thus, PBPK models are ideal candidates to simulate clinically important physical characteristics of patients, while simultaneously simulating biochemical and disease mechanisms on a systems level [61]. We (and many others in the systems pharmacology community) feel this is a critical unmet need, and an opportunity to connect basic science with clinical practice [62].

The performance of the effective complement model was impressive given its limited size. However, there are several questions that should be explored further. A logical progression for this work would be to expand the network to include the classical pathway and the formation of the membrane attack complex (MAC). However, time course measurements of MAC abundance (and MAC formation dynamics) are scarce, making the inclusion of MAC challenging. On the other hand, inclusion of classical pathway activation is straightforward. Liu et al., have shown cross-talk between the activation of the classical and lectin pathways through C reactive proteins (CRP) and L-ficolin (LF) under inflammation conditions [20]. Thus, inclusion of these species, in addition to a lumped activation term for the classical pathway should allow us to capture classical activation. Next, we should address the C3a time scale issue. We believe the C3a time scale was related to our choice of training data, how we modeled the tickover mechanism, and factor B and D limitation. Tickover was modeled as a first-order generation processes where C3wBb formation and activity was lumped into the AP C3 convertase. Thus, we skipped an important step which could strongly influence AP C3 convertase formation by slowing down the rate C3 cleavage into C3a and C3b. The model should be expanded to include the C3wBb intermediate, where C3wBb catalyzes C3 cleavage at a slow rate compared to normal AP or C4bC2as. We also assumed both factor B and factor D were not limiting, thereby artificially accelerating the rate of AP C3 convertase formation. This shortcoming could be addressed by including balances around factor B and D, and including these species in the appropriate kinetic rates. The C5a predictions also had an accelerated time scale. However, because the C5a time scale depended strongly upon C3 convertase formation, we can likely correct the C5 issues by fixing the rate of C3 cleavage. We should also consider including the C2-bypass pathway, which was not included in the model. The C2-bypass mediates lectin pathway activation, without the involvement of MASP-1/2; this pathway could be important for understanding the role of MASP-1/2 inhibitors on complement activation. Lastly, we have assumed that zymosan A differentially actives the lectin pathway, however, this may not be true. We should further explore the potential cross activation of all three branches by zymosan to determine the sensitivity of the model predictions to the assumption of differential activation.

## Author’s contributions

J.V directed the study. A.S developed the reduced order complement model and the parameter ensemble. A.S, W.D, R.L and M.M analyzed the model ensemble, and generated figures for the manuscript. The manuscript was prepared and edited for publication by A.S, W.D, M.M, R.L and J.V.

## Acknowledgements

We gratefully acknowledge the suggestions from the anonymous reviewers to improve this manuscript.

## Supplemental Information

**Table T1.** Complement protein initial conditions and associated species indexed used in the model. All unlisted proteins had an initial condition of zero.

| Protein | Index | Initial Condition | Units | Source |
| --- | --- | --- | --- | --- |
| C4 | 2 | 1.9 | $\mu\text{M}$ | [63] |
| C2 | 3 | 0.322 | $\mu\text{M}$ | [63] |
| C3 | 8 | 7.57 | $\mu\text{M}$ | [63] |
| C5 | 1 | 0.195 | $\mu\text{M}$ | [63] |
| Factor H | 17 | 2.23 | $\mu\text{M}$ | [63] |
| C4BP | 18 | 0.417 | $\mu\text{M}$ | [63] |

**Table T2.** Experimental measurements used in modeling training and validation from Morad and coworkers [34]. Two sets of C3a and C5a measurements were used for model training while three were used for validation. The information of the experimental data, its usage in this work and its location within the original publication is described below.

| Protein | zymosan ($\mu M$) | Figure Number | Model Utilization |
| --- | --- | --- | --- |
| C3a | 0 | Fig 2a | Training |
|  | 1.0 | Fig 2e | Training |
|  | 0.001 | Fig 2b | Prediction |
|  | 0.01 | Fig 2c | Prediction |
|  | 0.1 | Fig 2d | Prediction |
| C5a | 0 | Fig 3a(i) | Training |
|  | 1.0 | Fig 3c | Training |
|  | 0.001 | Fig 3a(ii) | Prediction |
|  | 0.01 | Fig 3a(iii) | Prediction |
|  | 0.1 | Fig 3b | Prediction |

**Table T3.** Model species table.

| Species | Index | Description |
| --- | --- | --- |
| Initiator2 | x[1] | Initiator complex 2, lectin pathway |
| C4 | x[2] | C4 protein |
| C2 | x[3] | C2 protein |
| C4a | x[4] | C4a protein |
| C4b | x[5] | C4b protein |
| C2a | x[6] | C2a protein |
| C2b | x[7] | C2b protein |
| C3 | x[8] | C3 protein |
| C3b | x[9] | C3b protein |
| C4bC2a | x[10] | Classical/Lectin C3 convertase |
| C3Convertase2 | x[11] | Alternate C3 convertase |
| C4bC2aC3b | x[12] | Classical C5 convertase |
| C5Convertase2 | x[13] | Alternate C5 convertase |
| C5 | x[14] | C5 protein |
| C5a | x[15] | C5a protein |
| C5b | x[16] | C5b protein |
| C4BP | x[17] | C4 binding protein |
| FactorH | x[18] | Factor H |
| C3a | x[19] | C3a protein |

